# Metagenomic DNA sequencing for the diagnosis of intraocular infections

**DOI:** 10.1101/109686

**Authors:** Thuy Doan, Nisha R. Acharya, Benjamin A. Pinsky, Malaya K. Sahoo, Eric D. Chow, Niaz Banaei, Indre Budvytiene, Vicky Cevallos, Lina Zhong, Zhaoxia Zhou, Thomas M. Lietman, Joseph L. DeRisi

**Affiliations:** Francis I. Proctor Foundation, University of California San Francisco, San Francisco, CA, USA; Department of Ophthalmology, University of California, San Francisco, CA, USA; Department of Pathology, Stanford University School of Medicine, Stanford, CA, USA; Department of Medicine, Division of Infectious Diseases and Geographic Medicine, Stanford University School of Medicine, Stanford, CA, USA; Department of Biochemistry and Biophysics, University of California, San Francisco, CA, USA; Stanford Health Care, Stanford, CA, USA; Chan Zuckerberg Biohub, San Francisco, CA, USA

**Author notes:** Corresponding Author:Joseph L. DeRisi, PhD, Department of Biochemistry and Biophysics, University of California, San Francisco, CA, USA., url http://derisilab.ucsf.edu.

## Abstract

**Purpose:** To compare the performance of unbiased high-throughput sequencing with pathogen directed PCR using DNA isolated from archived ocular fluid, approaches that are compatible with the current sample handling practice of ophthalmologists.

**Design:** Retrospective molecular study of banked vitreous samples.

**Methods:** We evaluated a metagenomic DNA sequencing-based approach (DNA-seq) using archived positive (*n* = 31) and negative (*n*=36) vitreous specimens as determined by reference pathogen-specific PCR assays (herpes simplex virus 1 and 2, cytomegalovirus, varicella-zoster virus, and *Toxoplasma gondii*). Pathogens were identified using a rapid computational pipeline to analyze the non-host sequences obtained from DNA-seq. Clinical samples were de-identified and laboratory personnel handling the samples and interpreting the data were masked.

**Results:** Metagenomic DNA sequencing detected 87% of positive reference samples. In the presumed negative reference samples, DNA-seq detected an additional 6 different pathogens in 8 samples (22% of negative samples) that were either not detected or not targeted with pathogen-specific PCR assays. Infectious agents identified only with DNA-seq were *Candida dubliniensis*, *Klebsiella pneumoniae*, human herpesvirus 6 (HHV-6), and human T-cell leukemia virus type 1 (HTLV-1). Discordant samples were independently verified in CLIA-certified laboratories. CMV sequences were compared against the antiviral mutation database and 3 of the samples were found to have mutations conferring ganciclovir resistance.

**Conclusions:** Metagenomic DNA sequencing was highly concordant with pathogen-directed PCRs. The unbiased nature of metagenomics DNA sequencing allowed an expanded scope of pathogen detection, including bacteria, fungal species, and viruses, resolving 22% of cases that had previously escaped detection by routine pathogen-specific PCRs. The detection of drug resistance mutations highlights the potential for unbiased sequencing to provide clinically actionable information beyond pathogen species detection.

## INTRODUCTION

The detection and validation of intraocular infections heavily rely on molecular diagnostics. Ocular samples can be challenging since as little as 100-300 µL of intraocular fluid can be safely obtained at any given time for diagnostic testing. The most widely available molecular diagnostic panel for infections in ophthalmology includes 4 separate pathogen-directed polymerase chain reactions (PCRs): cytomegalovirus (CMV), herpes simplex virus (HSV), varicella zoster virus (VZV), and *Toxoplasma gondii*. More than 50% of all presumed intraocular infections fail to have a pathogen identified.^1^

Metagenomic deep sequencing has the potential to improve diagnostic yield as it is inherently unbiased and hypothesis-free – it can theoretically detect all pathogens in a clinical sample and simultaneously provide genetic information that may be used for epidemiological investigations and antimicrobial susceptibility.^2,3^ Further, technical advances in metagenomic deep sequencing and bioinformatics have made it increasingly feasible for the translational application of this technology to the clinical setting (http://www.ciapm.org). Previously, we have demonstrated that unbiased RNA sequencing (RNA-seq) of intraocular fluid can detect fungi, parasites, DNA and RNA viruses in a proof-of-principle study for uveitis patients.^2^ One obvious drawback regarding RNA-seq is that optimal RNA sequencing results require proper specimen handling that includes either flash-freezing or immediately placing the specimen on dry ice. While commercial roomtemperature RNA-preservatives may address this issue, it is impractical in the near-term for the majority of the practicing ophthalmologists as the procedures to obtain intraocular fluids are routinely done in an outpatient setting. For pathogens with DNA genomes, metagenomic DNA sequencing (DNA-seq), can partially circumvent this challenge, as the quality of DNA can be more tolerant of specimen handling under ambient temperature, and the use of DNA makes available large retrospective sample collections for analysis. The purpose of this study was to compare the performance of metagenomic DNA sequencing against conventional pathogen-directed PCRs to diagnose intraocular infections.

## METHODS

### Samples

All archived vitreous samples were banked from January 2010 to December 2015. These vitreous samples were received at the F. I. Proctor Foundation for diagnostic PCR testing. Leftover samples were previously boiled and then stored at −80°C. These samples were de-identified using standard institutional procedures and randomized. Laboratory personnel involved in sample preparation, analysis, and confirmatory testing were masked to the reference PCR results until all analyses were completed.

### Pathogen-directed PCRs

The Proctor Foundation maintains a CLIA-certified laboratory that routinely performs HSV, VZV, CMV, and *Toxoplasma gondii* PCRs on intraocular fluid samples.^4–7^ Briefly, intraocular fluid samples are boiled for 10 minutes and 4 - 5 μL of the boiled vitreous is subjected to PCR with the respective pathogen-directed primers (see Table S1 in the supplementary materials). All pathogen-directed PCR assays used 5 μL of 10X PCR buffer (Sigma-Aldrich Corp., St. Louis, MO), 1.5 - 2.5 mM of MgCl_2_, 0.05 - 0.1 *m*M dNTP mix, 10pmol of each primer, and 0.05 – 0.08 U of REDTaq DNA polymerase (Sigma-Aldrich Corp., St. Louis, MO) in a total reaction volume of 50 - 100 μL. Amplified products were evaluated on 4% e-Gels (Thermo Fisher Scientific, Waltham, MA).

### Library Preparation and Sequencing

DNA was isolated from 50 μL of each vitreous sample using the DNeasy Blood & Tissue Kit (QIAGEN, Germantown, MD) per the manufacturer’s recommendations. The DNA was eluted in 9 μL of the kit elution buffer and stored at −20°C. DNA was quantified using the Qubit^®^ dsDNA HS Assay Kit (Thermo Fisher Scientific, Waltham, MA). 5 μL of the extracted DNA were used to prepare DNA-seq libraries using the Nextera XT DNA Library Preparation Kit (Illumina, San Diego, CA) according to the manufacturer’s recommendations and amplified with 12 PCR cycles. Library size and concentration were determined using the Blue Pippin (Sage Science, Beverly, MA) and Kapa Universal quantitative PCR kit (Kapa Biosystems, Woburn, MA), respectively. Samples were sequenced to an average depth of ~15 × 10^6^ reads/sample on an Illumina HiSeq 4000 instrument using 125 nucleotide (nt) paired-end sequencing.^8,9^ A water (“no-template”) control was included in each library preparation.

### Bioinformatics

Sequencing data were analyzed using a rapid computational pipeline developed by the DeRisi Laboratory to classify MDS reads and identify potential pathogens by comparison to the entire NCBI nucleotide (nt) reference database.^2,8^ Briefly, the pipeline consists of the following steps. First, an initial human-sequence removal step is accomplished by alignment of all paired-end reads to the human reference genome 38 (hg38) and the *Pan troglodytes* genome (panTro4, 2011, UCSC), using the Spliced Transcripts Alignment to a Reference (STAR) aligner (v2.5.1b).^10^ Unaligned reads were quality filtered using PriceSeqFilter^11^ with the “-rnf 90” and “-rqf 85 0.98” settings. Reads passing QC were then subjected to duplicate removal. The remaining reads that were at least 95% identical were compressed by cd-hit-dup (v4.6.1).^12^ Paired reads were then assessed for complexity by compression with the LZW algorithm.^13^ Read-pairs with a compression score less than 0.45 were subsequently removed. Next, a second phase of human removal was conducted using the very-sensitive-local mode of Bowtie2 (v2.2.4) with the same hg38 and PanTro4 reference as described above.^14^ Read pairs, in which both members remained unmapped, were then passed on to GSNAPL (v2015-12-31).^15^ At this step, read-pairs were aligned to the NCBI nt database (downloaded July 2015, indexed with k=16mers), and preprocessed to remove known repetitive sequence with RepeatMasker (vOpen-4.0) (www.repeatmasker.org). Finally, reads were aligned to the NCBI non-redundant nucleotide (nr) database (July 2015) using the Rapsearch2 algorithm.^16^ For the purpose of this study, human infectious agents detected by our pipeline were manually inspected using Bowtie2 to align against full-length reference genomes. Geneious (Biomatters, Ltd., Auckland, New Zealand) was used for visualization. No absolute or relative read counts were used to determine a call. Specifically, any infectious agent that had ≥ 2 non-overlapping reads to the reference pathogen genome was considered positive if it met both of the following criteria: (1) It is a pathogen known to be a cause of infectious uveitis and (2) reads from this pathogen were not present in the no-template (“water only”) control on the same run and library preparation.

### Confirmatory PCRs

Discrepant samples were evaluated at the CLIA-certified Stanford Clinical Microbiology and Clinical Virology laboratories, when possible. 50μL of ocular specimen, or all remaining volume if less than 50 μL, was diluted in Nuclease Free H_2_O (TekNova, Hollister, CA) to a total volume of 200 μL and total nucleic acids were extracted on the BioRobot EZ1 (QIAGEN, Germantown, MD) using the EZ1 virus mini kit 2.0 according to the manufacturer’s recommendations. CMV, HSV-1/HSV-2, and VZV were confirmed using *artus* real-time PCR analyte specific reagents on the Rotor-Gene Q instrument (both from QIAGEN, Germantown, MD). HHV-6 was evaluated using a laboratory developed real-time PCR protocol targeting the U66 gene (see supplemental methods). Bacterial and fungal agents were confirmed using PCR amplification and Sanger sequencing of the 16S rRNA gene, or the 28S rRNA gene D2 region and fungal rRNA internal transcribed spacer 2 (ITS2), respectively. *T.gondii* was tested using assays targeting both the B1 and Rep529 repetitive elements (see supplemental methods).

Detection of HTLV-1 was performed using the following primers^17^: HTLV1-F1 5’-ACAAAGTTAACCATGCTTATTATCAGC-3’ and HTLV1-R1 5’-ACACGTAGACTGGGTATCCGAA-3’. One-Step RT PCR with Platinum Taq master mix (Thermo Fisher Scientific, Waltham, MA) was used with the following amplification conditions: 55°C for 30 minutes, 40 cycles of 95°C for 30 seconds, 55°C for 30 seconds, and 72°C for 1 minute, with a final extension of 72°C for 5 minutes. The product was evaluated on a 2% agarose gel and visualized with Ethidium Bromide.

## RESULTS

### Metagenomic DNA sequencing of pathogen-directed PCR positive vitreous samples

To assess the sensitivity of DNA-seq, archived vitreous samples from 2010-2015, that were positive for HSV, VZV, HSV, and *T gondii*, were randomly selected (Figure 1). All samples were de-identified and randomized among PCR-negative samples. Of the 31 positive samples tested, 27 samples were identified correctly with DNA-seq (Table 1). 3 samples positive for *T gondii* and 1 sample positive for VZV by directed-PCR were not detected by DNA-seq. The positivity of these 4 samples was confirmed by directed PCRs at another CLIA-certified laboratory. The cycle threshold for VZV at the outside institution was 28.2, while the cycle thresholds for the three *T.gondii* samples ranged from 27-36.5. Therefore, the positive agreement between pathogens detected by directed-PCR and DNA-seq was 87%.

**Figure 1.**
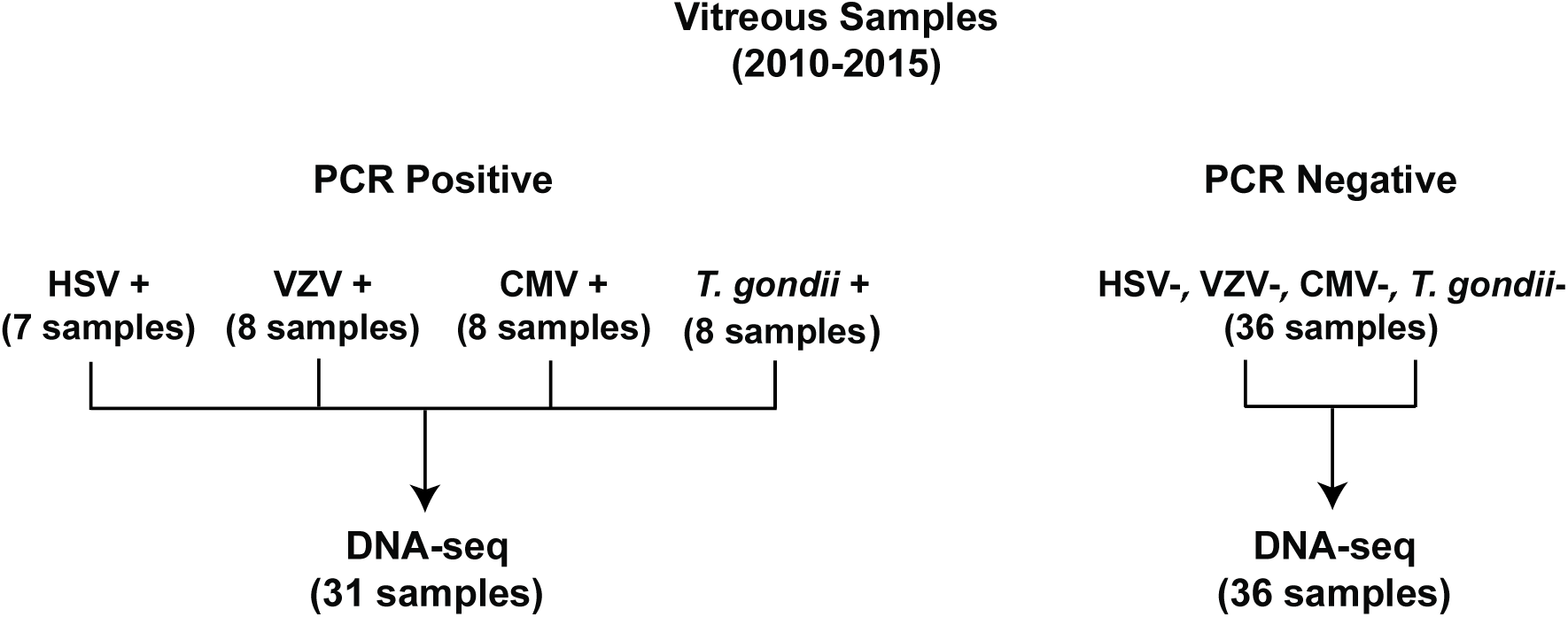
Study Profile. Pathogen-directed PCR positive and negative banked vitreous samples were subjected to DNA-seq.

**Table 1:** Result summary for reference PCR positive samples.

| Sample ID | Proctor PCR Results | DNA concentration (ng/μL) | DNA-seq Results | nt RM (sp) | Stanford PCR Results |
| --- | --- | --- | --- | --- | --- |
| 22 | CMV | 0.102 | CMV | 4.362 | NA |
| 28 | CMV | <0.05 | CMV | 537.635 | NA |
| 34 | CMV | 0.108 | CMV | 713.308 | NA |
| 40 | CMV | <0.05 | CMV | 21.663 | NA |
| 62 | CMV | 0.176 | CMV | 16.184 | NA |
| 65 | CMV | <0.05 | CMV | 286.589 | NA |
| 71 | CMV | 2.66 | CMV | 727.388 | NA |
| 74 | CMV | 0.134 | CMV | 982.894 | NA |
| 42 | HSV-1 | 1.06 | HSV-1 | 10.443 | NA |
| 17 | HSV-2 | 0.956 | HSV-2 | 3.751 | NA |
| 23 | HSV-2 | 0.682 | HSV-2 | 100.162 | NA |
| 36 | HSV-2 | 0.164 | HSV-2 | 40.482 | NA |
| 38 | HSV-2 | 0.100 | HSV-2 | 0.427 | NA |
| 47 | HSV-2 | 2.32 | HSV-2 | 49.161 | NA |
| 64 | HSV-2 | <0.05 | HSV-2 | 0.109 | NA |
| 15 | *T. gondii* | 1.90 | *T. gondii* | 34.936 | NA |
| 25* | *T. gondii* | 0.140 | Negative | NA | *T. gondii* |
| 30 | *T. gondii* | <0.05 | *T. gondii* | 49.609 | NA |
| 41 | *T. gondii* | 0.134 | *T. gondii* | 0.171 | NA |
| 50 | *T. gondii* | <0.05 | *T. gondii* | 0.653 | NA |
| 54* | *T. gondii* | 1.24 | Negative | NA | *T. gondii* |
| 57 | *T. gondii* | 0.180 | *T. gondii* | 256.063 | NA |
| 60* | *T. gondii* | 0.178 | Negative | NA | *T. gondii* |
| 14 | VZV | 0.538 | VZV | 0.41 | NA |
| 27 | VZV | <0.05 | VZV | 19.32 | NA |
| 29 | VZV | 4.16 | VZV | 8.395 | NA |
| 32 | VZV | 0.160 | VZV | 125.516 | NA |
| 43 | VZV | 3.92 | VZV | 2.585 | NA |
| 44* | VZV | 0.126 | Negative | NA | VZV |
| 48 | VZV | 1.01 | VZV | 1.214 | NA |
| 66 | VZV | 0.128 | VZV | 1.181 | NA |

\*Discordant sample
NA: Not applicable

### Metagenomic DNA sequencing of PCR negative vitreous samples

Thirty-six archived vitreous samples that tested negative by all Proctor pathogen-directed PCRs (CMV, VZV, HSV, and *T. gondii*) were randomly selected for DNA-seq. 28 out of 36 samples yielded no additional pathogen (Table 2), while 8 samples (22%) resulted in 6 additional pathogens either not detected or not tested with pathogen-directed PCRs. Those organisms included CMV (n = 2), HHV-6 (n = 2), HSV-2 (n = 1), HTLV-1 (n = 1), *Klebsiella pneumoniae* (n = 1), and *Candida dubliniensis* (n = 1). All of these organisms are known to cause infectious uveitis. Confirmatory testing of these pathogens, except HTLV-1, was performed at another CLIA-certified laboratory using assays validated for diagnostic testing. The results for HHV-6, *Klebsiella pneumoniae*, and *Candida dubliniensis* were confirmed. However, the presence of CMV DNA was confirmed by directed-PCR in only one of the two samples (Ct = 21.39). The sample not confirmed by directed-PCR had only 2 CMV reads by DNA-seq. Similarly, HSV-2 was not confirmed by PCR and again DNA-seq provided only 2 HSV-2 reads. Here, it was unclear if DNA-seq can achieve higher sensitivity for CMV and HSV-2 compared to directed PCR, or whether these were false positive DNA-seq results. It should be noted that DNA-seq detected 100% of all samples that tested positive for CMV and HSV-2 by PCRs. The vitreous sample positive for HTLV-1 by DNA-seq was subjected to HTLV-1 directed PCR with primers targeting the *tax* gene.^17^ The PCR product was sequenced (Elim Biosciences, Hayward, CA), which confirmed the presence of HTLV-1 DNA in this vitreous sample.

**Table 2:** Result summary for reference PCR pan-negative samples.

| Sample ID | Proctor PCR Results | DNA concentration (ng/μL) | DNA-seq Results | nt RM (sp) | Stanford PCR Results |
| --- | --- | --- | --- | --- | --- |
| 4 | Pan-Negative | <0.05 | Negative | NA | NA |
| 5 | Pan-Negative | <0.05 | Negative | NA | NA |
| 11 | Pan-Negative | <0.05 | Negative | NA | NA |
| 12 | Pan-Negative | 5.06 | Negative | NA | NA |
| 13 | Pan-Negative | <0.05 | Negative | NA | NA |
| 16 | Pan-Negative | <0.05 | Negative | NA | NA |
| 18* | Pan-Negative | <0.05 | *Candida dubliniensis* | 33.99 | *Candida dubliniensis* |
| 21 | Pan-Negative | 0.170 | Negative | NA | NA |
| 24 | Pan-Negative | <0.05 | Negative | NA | NA |
| 26 | Pan-Negative | <0.05 | Negative | NA | NA |
| 31 | Pan-Negative | 0.378 | Negative | NA | NA |
| 33 | Pan-Negative | <0.05 | Negative | NA | NA |
| 35 | Pan-Negative | <0.05 | Negative | NA | NA |
| 37 | Pan-Negative | <0.05 | Negative | NA | NA |
| 39* | Pan-Negative | <0.05 | HSV-2 | 0.064 | NDET |
| 45 | Pan-Negative | <0.05 | Negative | NA | NA |
| 46 | Pan-Negative | <0.05 | Negative | NA | NA |
| 49 | Pan-Negative | <0.05 | Negative | NA | NA |
| 51 | Pan-Negative | 0.692 | Negative | NA | NA |
| 52 | Pan-Negative | <0.05 | Negative | NA | NA |
| 53* | Pan-Negative | <0.05 | CMV | 0.045 | NDET |
| 55* | Pan-Negative | <0.05 | CMV | 1123.4 | CMV |
| 56 | Pan-Negative | <0.05 | Negative | NA | NA |
| 58 | Pan-Negative | <0.05 | Negative | NA | NA |
| 59 | Pan-Negative | 0.388 | Negative | NA | NA |
| 61 | Pan-Negative | 0.180 | Negative | NA | NA |
| 63* | Pan-Negative | <0.05 | HTLV-1 | 0.111 | HTLV-1 |
| 67 | Pan-Negative | 0.326 | Negative | NA | NA |
| 68* | Pan-Negative | <0.05 | HHV-6 | 2.79 | HHV-6 |
| 69 | Pan-Negative | <0.05 | Negative | NA | NA |
| 70* | Pan-Negative | 0.554 | HHV-6 | 2.583 | HHV-6 |
| 72 | Pan-Negative | 0.126 | Negative | NA | NA |
| 73 | Pan-Negative | <0.05 | Negative | NA | NA |
| 75* | Pan-Negative | 1.43 | *Klebsiella pneumoniae* | 9446.854 | *Klebsiella pneumoniae* |
| 76 | Pan-Negative | <0.05 | Negative | NA | NA |
| 77 | Pan-Negative | <0.05 | Negative | NA | NA |

\*Discordant sample
NA: Not applicable
NDET: Not detected

### Characteristics of metagenomic DNA sequencing

Non-host sequences represented a small fraction (0.7% ± 2.5%, mean ± SD) of the total number of reads consistent with prior metagenomic deep sequencing of clinical samples.^2,3^ The number of reads for the pathogen identified ranged from 2 to 166,691 reads or 0.045 to 9446 matched read pairs per million read pairs (rM) at the species level based on nucleotide alignment (Figure 2A). Of the samples considered pathogen positive by DNA-seq, the starting DNA concentration was significantly higher than samples that were considered pathogen negative (0.63 ± 1.05 vs 0.28 ± 0.95 ng/μL, mean ± SD, *p =* 0.01, Figure 2B), although a low DNA concentration did not exclude the identification of a pathogen or vice versa. In this study, the vitreous sample with the highest DNA concentration, 5.06 ng/μL, was not positive for a pathogen, suggesting that inflammatory cell infiltration may be robust even in the absence of infection.

**Figure 2.**
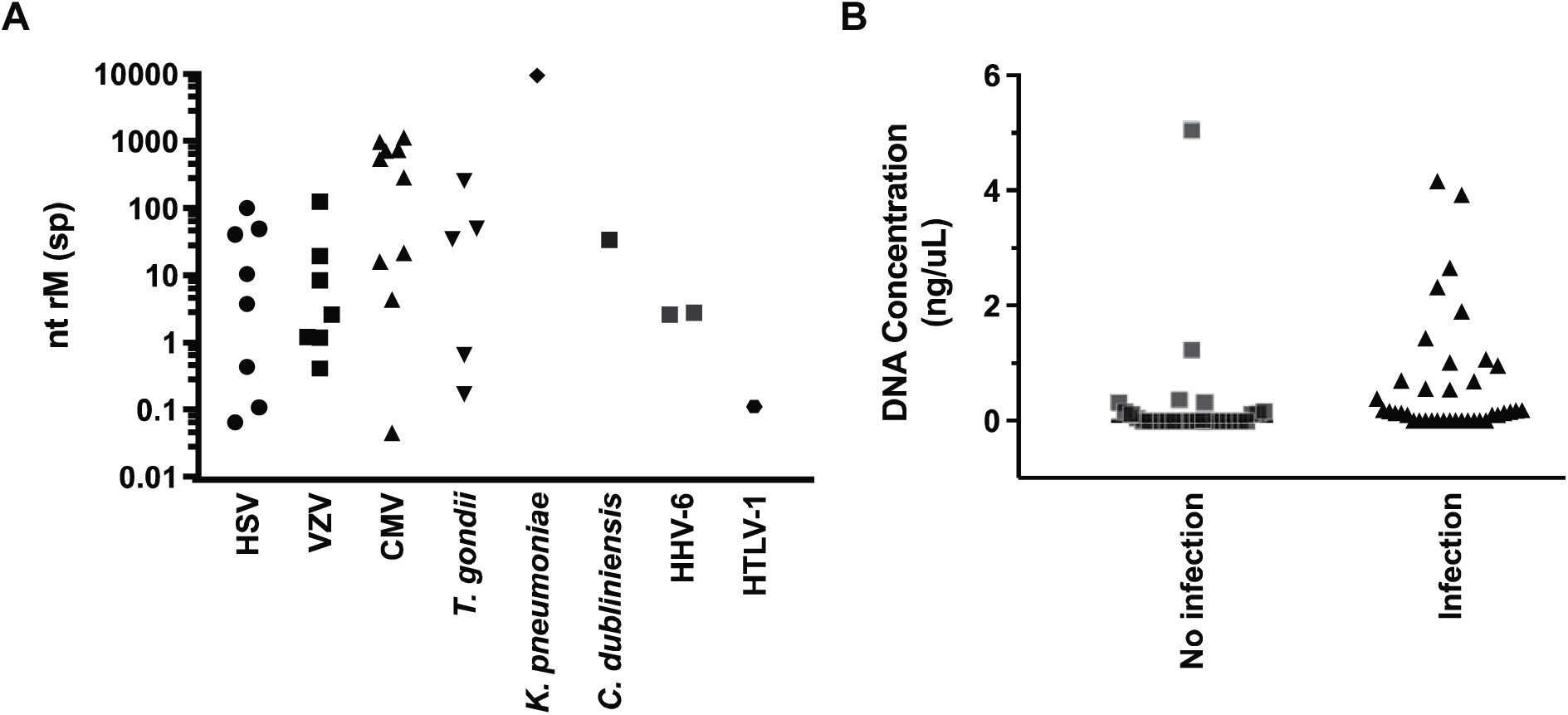
Characteristics of metagenomic DNA sequencing. (A) Normalized pathogen readcounts as detected by DNA-seq. Organisms in each sample are plotted as a function of matched read pairs per million read pairs (rM) at the species level based on nucleotide (nt) alignment. (B) Samples with infectious agents as detected by DNA-seq had significantly higher DNA concentration (0.63 ± 1.05 vs0.28 ± 0.95ng/μL, mean ± SD, *p* = 0.01, Mann-Whitney test) compared to negative samples

### Antiviral drug mutation analysis with metagenomic DNA sequencing

An advantage of metagenomic deep sequencing is the ability to apply sequence information to infer the phenotypic behavior of the identified pathogen. As a proof-of-concept, we compared samples in which CMV sequences were adequately recovered for the UL54 and UL97 genes, coding for the DNA polymerase and phosphotransferase respectively, and compared to a CMV antiviral drug resistance database developed at Stanford University ^18,19^. Of the 7 samples analyzed, 3 had mutations in UL97 that confer ganciclovir and valganciclovir resistance. Two samples were found to have C592G mutations, and 1 sample had both C592G and L595S mutations (Table 3). No drug resistance mutations were identified in UL54.

**Table 3:** CMV drug resistance mutations detected in banked vitreous samples. CMV, cytomegalovirus; UL97, phosphor transferase; UL54, DNA polymerase.

| Sample ID | Gene name | Drug resistance mutations | Predicted drug resistance pattern |
| --- | --- | --- | --- |
| 28 | UL97 | None |  |
|  | UL54 | None |  |
| 34 | UL97 | None |  |
|  | UL54 | None |  |
| 40 | UL97 | None |  |
|  | UL54 | None |  |
| 62 | UL97 | None |  |
|  | UL54 | None |  |
| 65 | UL97 | C592G | Ganciclovir resistant |
|  | UL54 | None |  |
| 71 | UL97 | C592G | Ganciclovir resistant |
| 74 | UL97 | C592G, L595S | Ganciclovir resistant |

## DISCUSSION

In this study, we showed that metagenomic DNA sequencing can detect most infections found by pathogen-directed PCR, with a positive agreement rate of 87% in intraocular fluid clinical samples. Further, DNA-seq detected a pathogen in an additional 22% of the samples that were either missed or cannot be detected with the available pathogen-directed PCR panel validated for ocular samples. These results highlight the robustness of DNA-seq given that different DNA extraction protocols were used and the samples that underwent DNA-seq were subjected to multiple additional freeze-thaw cycles compared to the original PCR testing.

DNA-seq correctly identified all HSV and CMV positive reference clinical samples. For the 4 pathogen-directed PCRs, DNA-seq was more sensitive than CMV-directed PCR as it identified an additional 2 samples with CMV DNA, but was less sensitive than *T. gondii*-directed PCR (62.5%, 5 out 8 samples). *T. gondii* is a widespread protozoan parasite with a 63 megabase-pair (Mbp) genome that includes 35 B1 gene repeats.^20^ The Proctor assay is a nested-PCR targeting the B1 gene with a limit of detection of ~5 *T. gondii* genomes/µL or 20 genomes/reaction. The lower sensitivity of DNA-seq for *T. gondii* compared to the Proctor assay is surprising given that this shotgun approach can theoretically detect the entire 63 Mbp genome of *T. gondii*. However, the pre-analytical techniques employed in this study may not be optimized for the detection of *T.gondii.* Future experiments comparing different intraocular specimen types, including fresh and frozen samples, as well as evaluation of extraction methods and library preparation protocols, may be necessary to improve *T. gondii* detection without compromising the sensitivity of other pathogens. Furthermore, it is possible that RNA-seq may provide better *T. gondii* sensitivity in intraocular fluid samples as prior work showed that this method can detect RNA from this organism with high genome coverage.^2^ While RNA was not available for evaluation from the samples used in this study these data suggest that a combined approach should be assessed, whereby both the RNA and DNA fractions are independently examined by metagenomic sequencing.

The benefit of target agnostic nature metagenomic sequencing is apparent here when applied to clinical samples that have tested negative with available pathogen-directed PCR assays. Of the 36 samples that tested negative at Proctor, 8 samples were positive for an intraocular pathogen with DNA-seq. Therefore, one in five patients with negative conventional testing had an intraocular infection that could have been more effectively treated. The number of missed pathogens is likely an underestimation, as DNA-seq cannot detect RNA viruses, such as rubella. These data suggests a practical diagnostic decision tree whereby samples negative by routine PCR are then advanced to metagenomic DNA and RNA sequencing.

An advantage of metagenomic sequencing is the ability to capture a large percentage of the organism’s genome, if not its entirety. Sequencing data can provide useful clinical information such as drug resistance. For 70% of the samples that were CMV positive by DNA-seq, coverage in the UL54 and UL97 genes allowed assessment for antiviral drug resistance mutations. We found that 43% of the samples (3 out of 7) analyzed had ganciclovir resistance mutations in the phosphotransferase gene (UL97). The two mutations identified were two of the most common CMV drug resistance mutations,^21^ indicating that these mutations were unlikely to represent sequencing errors. A caveat in interpreting these results is that the authors do not have information on the medication history of the patients identified to have ganciclovir resistant CMV infections.

In summary, we have shown that metagenomic DNA sequencing can detect a wide range of pathogens including bacteria, fungi, parasites, and DNA viruses in intraocular samples, concurrently provide drug resistance information, and improve diagnostic yield for samples that have failed conventional diagnostics. This approach will not only complement the current diagnostic paradigm in ophthalmology but also allow for a more comprehensive characterization of the etiology of infectious uveitis.

## ACKNOWLEDGMENTS

We thank Derek Bogdanoff in the UCSF Center for Advanced Technology for his expertise and assistance operating the Illumina sequencer. We thank Drs. Chaz Langelier and Michael Wilson for helpful discussions.

## REFERENCES

1. Sugita S, Ogawa M, Shimizu N, et al.Use of a comprehensive polymerase chain reaction system for diagnosis of ocular infectious diseases. Ophthalmology 2013; 120(9): 1761–8.

2. Doan T, Wilson MR, Crawford ED, et al.Illuminating uveitis: metagenomic deep sequencing identifies common and rare pathogens. Genome medicine 2016; 8(1): 90.

3. Graf EH, Simmon KE, Tardif KD, et al. Unbiased Detection of Respiratory Viruses by Use of RNA Sequencing-Based Metagenomics: a Systematic Comparison to a Commercial PCR Panel. Journal of clinical microbiology 2016; 54(4): 1000–7.

4. Cunningham ETJr., Short GA, Irvine AR, Duker JS, Margolis TP. Acquired immunodeficiency syndrome–associated herpes simplex virus retinitis. Clinical description and use of a polymerase chain reaction–based assay as a diagnostic tool. Arch Ophthalmol 1996; 114 (7): 834–40.

5. Short GA, Margolis TP, Kuppermann BD, Irvine AR, Martin DF, Chandler D. A polymerase chain reaction-based assay for diagnosing varicella-zoster virus retinitis in patients with acquired immunodeficiency syndrome. American journal of ophthalmology 1997; 123(2): 157–64.

6. McCann JD, Margolis TP, Wong MG, et al.A sensitive and specific polymerase chain reaction-based assay for the diagnosis of cytomegalovirus retinitis. American journal of ophthalmology 1995; 120(2): 219–26.

7. Grigg ME, Ganatra J, Boothroyd JC, Margolis TP. Unusual abundance of atypical strains associated with human ocular toxoplasmosis. The Journal of infectious diseases 2001; 184(5): 633–9.

8. Wilson MR, Shanbhag NM, Reid MJ, et al.Diagnosing Balamuthia mandrillaris Encephalitis With Metagenomic Deep Sequencing. Annals of neurology 2015.

9. Wilson MR, Naccache SN, Samayoa E, et al. Actionable diagnosis of neuroleptospirosis by next-generation sequencing. N Engl J Med 2014; 370(25): 2408–17.

10. Dobin A, Davis CA, Schlesinger F, et al. STAR: ultrafast universal RNA-seq aligner. Bioinformatics (Oxford, England) 2013; 29(1): 15–21.

11. Ruby JG, Bellare P, Derisi JL. PRICE: software for the targeted assembly of components of (Meta) genomic sequence data. G3 (Bethesda, Md) 2013; 3(5): 865–80.

12. Fu L, Niu B, Zhu Z, Wu S, Li W. CD-HIT: accelerated for clustering the next-generation sequencing data. Bioinformatics 2012; 28(23): 3150–2.

13. Ziv J, Lempel A. A universal algorithm for sequential data compression. IEEE Transactions on Information Theory 1977; 23(3): 337–43.

14. Langmead B, Salzberg SL., Fast gapped-read alignment with Bowtie 2. Nature methods 2012; 9(4): 357–9.

15. Wu TD, Nacu S. Fast and SNP-tolerant detection of complex variants and splicing in short reads. Bioinformatics (Oxford, England) 2010; 26(7): 873–81.

16. Zhao Y, Tang H, Ye Y. RAPSearch2: a fast and memory-efficient protein similarity search tool for next-generation sequencing data. Bioinformatics (Oxford, England) 2012; 28(1): 125–6.

17. Brunetto GS, Massoud R, Leibovitch EC, et al.Digital droplet PCR (ddPCR) for the precise quantification of human T-lymphotropic virus 1 proviral loads in peripheral blood and cerebrospinal fluid of HAM/TSP patients and identification of viral mutations. Journal of neurovirology 2014; 20(4): 341–51.

18. Sahoo MK, Lefterova MI, Yamamoto F, et al.Detection of cytomegalovirus drug resistance mutations by next-generation sequencing. J Clin Microbiol 2013; 51(11): 3700–10.

19. Chou S, Ercolani RJ, Sahoo MK, Lefterova MI, Strasfeld LM, Pinsky BA. Improved detection of emerging drug-resistant mutant cytomegalovirus subpopulations by deep sequencing. Antimicrob Agents Chemother 2014; 58(8): 4697–702.

20. Burg JL, Grover CM, Pouletty P, Boothroyd JC. Direct and sensitive detection of a pathogenic protozoan, Toxoplasma gondii, by polymerase chain reaction. Journal of clinical microbiology 1989; 27(8): 1787–92.

21. Lurain NS, Chou S.Antiviral drug resistance of human cytomegalovirus. Clinical microbiology reviews 2010; 23(4): 689–712.

22. Fekkar A, Bodaghi B, Touafek F, Le Hoang P, Mazier D, Paris L. Comparison of immunoblotting, calculation of the Goldmann-Witmer coefficient, and real-time PCR using aqueous humor samples for diagnosis of ocular toxoplasmosis. Journal of clinical microbiology 2008; 46(6): 1965–7.

